# MitoFinder: efficient automated large-scale extraction of mitogenomic data in target enrichment phylogenomics

**DOI:** 10.1101/685412

**Authors:** Rémi Allio, Alex Schomaker-Bastos, Jonathan Romiguier, Francisco Prosdocimi, Benoit Nabholz, Frédéric Delsuc

## Abstract

Thanks to the development of high-throughput sequencing technologies, target enrichment sequencing of nuclear ultraconserved DNA elements (UCEs) now allows routinely inferring phylogenetic relationships from thousands of genomic markers. Recently, it has been shown that mitochondrial DNA (mtDNA) is frequently sequenced alongside the targeted loci in such capture experiments. Despite its broad evolutionary interest, mtDNA is rarely assembled and used in conjunction with nuclear markers in capture-based studies. Here, we developed MitoFinder, a user-friendly bioinformatic pipeline, to efficiently assemble and annotate mitogenomic data from hundreds of UCE libraries. As a case study, we used ants (Formicidae) for which 501 UCE libraries have been sequenced whereas only 29 mitogenomes are available. We compared the efficiency of four different assemblers (IDBA-UD, MEGAHIT, MetaSPAdes, and Trinity) for assembling both UCE and mtDNA loci. Using MitoFinder, we show that metagenomic assemblers, in particular MetaSPAdes, are well suited to assemble both UCEs and mtDNA. Mitogenomic signal was successfully extracted from all 501 UCE libraries allowing confirming species identification using COI barcoding. Moreover, our automated procedure retrieved 296 cases in which the mitochondrial genome was assembled in a single contig, thus increasing the number of available ant mitogenomes by an order of magnitude. By leveraging the power of metagenomic assemblers, MitoFinder provides an efficient tool to extract complementary mitogenomic data from UCE libraries, allowing testing for potential mito-nuclear discordance. Our approach is potentially applicable to other sequence capture methods, transcriptomic data, and whole genome shotgun sequencing in diverse taxa.

## Introduction

Next generation phylogenomics in which phylogenetic relationships are inferred from thousands of genomic markers gathered through high-throughput sequencing (HTS) is on the rise. More specifically, targeted enrichment or DNA sequence capture methods are becoming the gold standard in phylogenetic analyses because they allow subsampling the genome efficiently at reduced cost (Lemmon & Lemmon, 2013; McCormack, Hird, Zellmer, Carstens, & Brumfield 2013a). The field has witnessed the rapid parallel development of exon capture from transcriptome-derived baits (Bi *et al.* 2012), anchored hybrid enrichment techniques (Lemmon, Emme, & Lemmon 2012), and the capture of ultraconserved DNA elements (UCEs; McCormack, Hird, Zellmer, Carstens, & Brumfield 2013b). All hybridization capture methods target a particular portion of the genome corresponding to the defined probes plus flanking regions. Prior knowledge is required to generate sequence capture probes, but ethanol preserved tissues, old DNA extractions, and museum specimens can be successfully sequenced (Faircloth *et al.* 2012; Guschanski *et al.* 2013; Blaimer *et al.* 2015). The first UCEs were identified by Bejerano *et al.* (2004) in the human genome and have been shown to be conserved in mammals, birds, and even ray-finned fish. Thanks to their large-scale sequence conservation, UCEs are particularly well-suited for sequence capture experiments and have become popular for phylogenomic reconstruction of diverse animals groups (Guschanski *et al.* 2013; Blaimer *et al.* 2015; Esselstyn, Oliveros, Swanson, & Faircloth 2017). Initially restricted to a few vertebrate groups such as mammals (McCormack *et al.* 2012) and birds (McCormack *et al.* 2013a), new UCE probe sets have been designed to target thousands of loci in arthropods such as hymenopterans (Blaimer *et al.* 2015; Branstetter *et al.* 2017a; Faircloth, Branstetter, White, & Brady 2015), coleopterans (Baca, Alexander, Gustafson, & Short 2017), and arachnids (Starrett *et al.* 2017).

It has been shown that complete mitochondrial genomes could be retrieved as by-products of sequence capture/enrichment experiments such as whole exome capture in human (Picardi & Pesole, 2012). Indeed, mitogenomes can in most cases be assembled from off-target sequences of UCE capture libraries in amniotes (do Amaral *et al.* 2015). Despite its well-acknowledged limitations (Galtier, Nabholz, Glémin, & Hurst 2009), mitochondrial DNA (mtDNA) remains a marker of choice for phylogenetic inference (e.g. Hassanin *et al.* 2012), for species identification or delimitation through barcoding (e.g. Coissac *et al.* 2016), and to reveal potential cases of mito-nuclear discordance resulting from introgression and/or hybridization events (e.g. Zarza et al. 2016, 2018; Grummer, Morando, Avila, Sites Jr, & Leaché 2018). MtDNA could also be used to taxonomically validate the specimens sequenced for UCEs using COI barcoding (Ratnasingham & Hebert, 2007) and to control for potential cross-contaminations in HTS experiments (Ballenghien, Faivre, & Galtier 2017). In practice, the few studies that have extracted mtDNA signal from UCEs (*e.g.* Meiklejohn *et al.* 2014; Pie *et al.* 2017; Wang, Hosner, Liang, Braun, & Kimball 2017, Zarza *et al.* 2018) and anchored phylogenomics (Caparroz *et al.* 2018) have done so manually for only few taxa. Most studies assembling mitogenomes from UCE libraries have used contigs produced by the Trinity RNAseq assembler (Grabherr *et al.* 2011) as part of the PHYLUCE pipeline (Faircloth, 2016), which was specifically designed to extract UCE loci. Indeed, RNAseq assemblers such as Trinity allow dealing with the uneven coverage of target reads in sequence-capture libraries, but also multi-copy genes such as the ribosomal RNA cluster, and organelles (chloroplasts and mitochondria). However, this strategy is likely not scaling well with hundreds of taxa because of the high computational demand required by Trinity. Metagenomic assemblers could provide a powerful alternative because they have been designed for an efficient *de novo* assembly of complex read populations by explicitly dealing with uneven read coverage and are computationally and memory efficient. Comparisons based on empirical bulk datasets of known composition (Vollmers, Wiegand, & Kaster 2017) have identified IDBA-UD (Peng, Leung, Yiu, & Chin 2012), MEGAHIT (Li *et al.* 2016), and MetaSPAdes (Nurk, Meleshko, Korobeynikov, & Pevzner 2017) as the most efficient current metagenomic assemblers.

As a case study, we focused on ants (Hymenoptera: Formicidae) for which a only 29 mitogenomes were available on GenBank compared to 501 UCE captured libraries as of March 29^th^, 2018 (**Appendix S1**). This contrasts sharply with the other most speciose group of social insects, termites (Isoptera), for which almost 500 reference mitogenomes have been produced (Bourguignon *et al.* 2017) and no UCE study has been conducted so far. Sequencing and assembling difficulties stemming from both the AT-rich composition (Foster, Jermiin, & Hickey 1997) and a high rate of mitochondrial genome rearrangements in hymenopteran (Dowton, Castro, & Austin 2002) might explain the limited number of mitogenomes currently available for ants. It is only recently that a few ant mitogenomes have been assembled out from UCE data (Ströher *et al.* 2017; Meza-Lázaro, Poteaux, Bayona-Vásquez, Branstetter, & Zaldívar-Riverón 2018; Vieira & Prosdocimi, 2019). Here, we built a pipeline called MitoFinder designed to automatically assemble and extract mitogenomic data from raw UCE capture libraries. Using publicly available UCE libraries for 501 ants, we show that complementary mitochondrial phylogenetic signal can be efficiently extracted using metagenome assemblers along with targeted UCE loci.

## Materials and methods

### Data acquisition

We used UCE raw sequencing data for 501 ants produced in 10 phylogenomic studies (Blaimer *et al.* 2015; Faircloth *et al.* 2015; Blaimer *et al.* 2016; Branstetter *et al.* 2017a,b,c; Jesovnik *et al.* 2017; Pierce *et al.* 2017; Prebus *et al.* 2017; Ward & Branstetter 2017). This dataset includes representatives of 15 of 16 subfamilies (Ward 2014) and 30 tribes. Raw sequence reads were downloaded from the NCBI Short Read Archive (SRA) on March 29^th^, 2018 (**Appendix S1**). For the 501 ant UCE libraries, raw reads were cleaned with Trimmomatic v0.36 (Bolger, Lohse, & Usadel 2014) using the following parameters: LEADING:3 TRAILING:3 SLIDINGWINDOW:4:15 MINLEN:50. A reference database with the 29 complete mitochondrial genomes available for ants on GenBank at the time was constructed.

### Mitogenomic data extraction with MitoFinder

To extract mitogenomic data from UCE libraries, we developed a dedicated bioinformatic pipeline called MitoFinder (**Fig. 1**). This pipeline was designed to assemble sequencing reads from target enrichment libraries, assemble, extract, and annotate mitochondrial contigs. To evaluate the impact of assembler choice, contigs were assembled with IDBA-UD v1.1.1, MEGAHIT v1.1.3, and MetaSPAdes v3.13.0 within MitoFinder, and with Trinity v2.1.1 within PHYLUCE using default parameters. Mitochondrial contigs were then identified by similarity search using blastn with e-value ≥ 1e-06 against our ant reference mitogenomic database. Each detected mitochondrial contig was then annotated with tblastx for protein-coding genes (CDS) and blastn for both 16S and 12S taking advantage of the geneChecker module of mitoMaker (Schomaker-Bastos & Prosdocimi, 2018) that we incorporated in MitoFinder. Finally, we used ARWEN v1.2 (Laslett & Canbäck, 2007) to detect and annotate tRNA genes.

**Figure 1.**
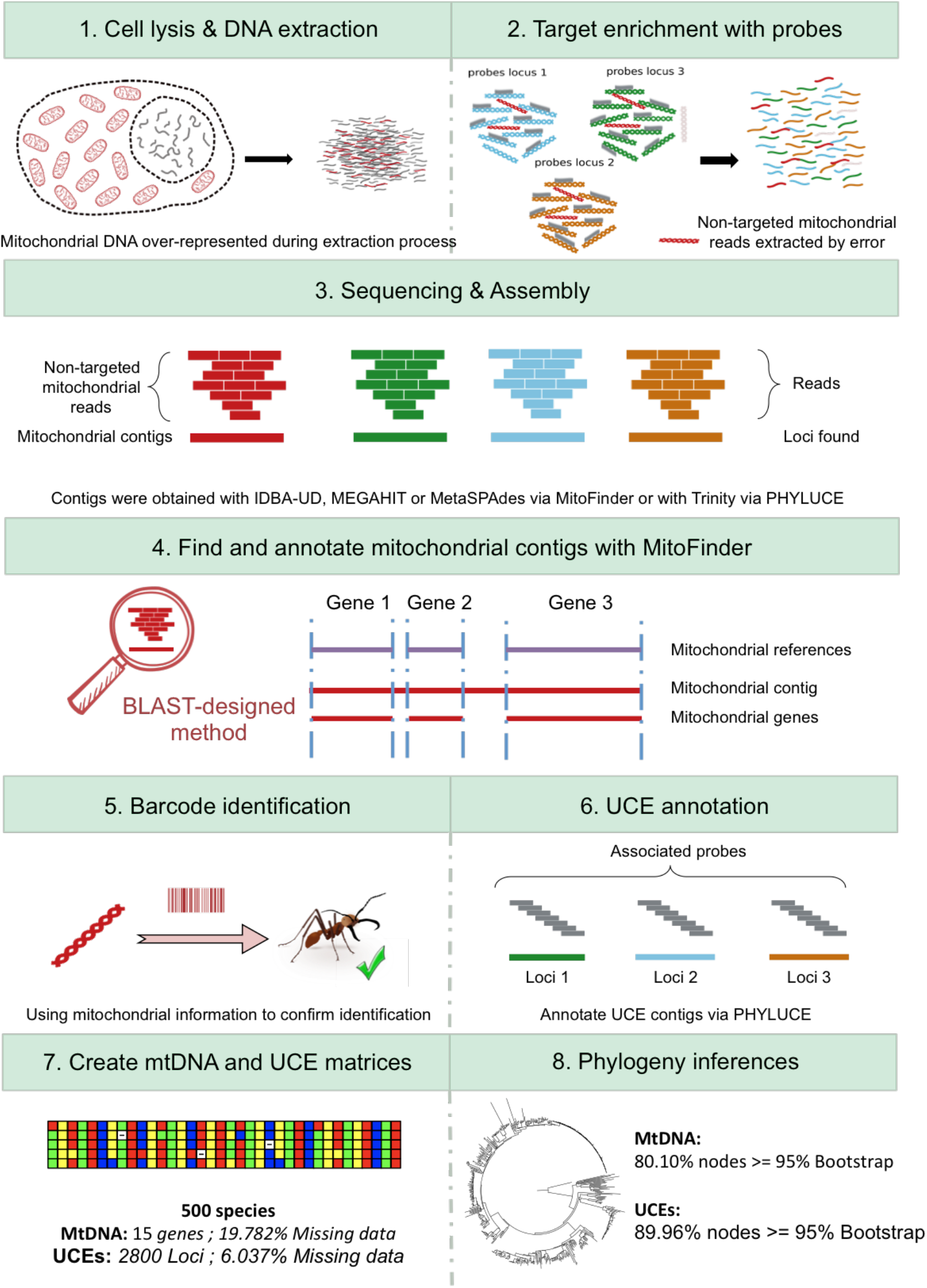
Conceptualization of the pipeline used to assemble and extract UCE and mitochondrial signal from ultraconserved element sequencing data.

Considering possible rearrangements in ant mitogenomes, each mitochondrial CDS was first aligned with MAFFT v7.271 (Katoh & Standley, 2013) algorithm FFT-NS-2 with option --*adjustdirection*. Then, to take into account potential frameshifts and stop codons, mitochondrial CDSs were re-aligned with MACSE v2.03 (Ranwez *et al.* 2018) with option - prog alignSequences, which produces both nucleotide and amino acid alignments. To improve alignment accuracy and reduce calculation time, we used sequences from available mitogenomes as references for each CDS (option *-seq_lr*). Sequences with internal stop codons were excluded to remove incorrectly annotated fragments potentially corresponding to nuclear mitochondrial DNA segments (NUMTs) in each protein-coding gene alignment. Then, individual gene alignments were eye-checked to manually remove remaining aberant sequences. Finally, a nucleotide supermatrix was created by concatenating protein-coding and ribosomal RNA genes. Considering mitochondrial signal saturation with high divergence, an amino acid supermatrix with the 13 mitochondrial CDSs was also assembled.

### DNA barcoding

To verify species identification of the 501 ant UCE libraries, COI sequences extracted by MitoFinder using MetaSPAdes (mtDNA recovered for all species) were compared with Species Level Barcode Records (3,328,881 COI sequences including more than 100,000 ants) through the identification server of the Barcode Of Life Data System v4 (Ratnasingham & Hebert, 2007). The same COI sequences were also compared against the NCBI nucleotide database using Megablast with default parameters.

### Assembly of UCEs

As recommended by Faircloth (2016), we first relied on Trinity to assemble UCE contigs using the *phyluce_assembly_assemblo_trinity* module of PHYLUCE. To assess the impact of assembler choice on UCE retrieval, we also used the assemblies obtained with IDBA-UD, MEGAHIT, and MetaSPAdes as implemented in MitoFinder. PHYLUCE scripts *phyluce_assembly_get_match_counts* and *phyluce_assembly_get_fastas_from_match_counts* were used to match contigs obtained for each sample to the bait set targeting 2590 UCE loci for Hymenoptera (Branstetter *et al.* 2017b). The resulting alignments were then cleaned using Gblocks (Castresana 2000) with the *phyluce_align_get_gblocks_trimmed_alignments_from_untrimmed* script. Finally, loci found in at least 75% of species were selected to create the four corresponding UCE supermatrices using the *phyluce_align_get_only_loci_with_min_taxa* script.

### Phylogenetic analyses

Phylogenetic relationships of ants were inferred from a total of 16 different supermatrices corresponding to the four supermatrices constructed from contigs obtained with each of the four assemblers (IDBA-UD, MEGAHIT, MetaSPAdes, and Trinity). The four supermatrices are as follows: (i) a UCE nucleotide supermatrix built from the concatenation of UCE loci retrieved for at least 75% of species, (ii) a mitochondrial nucleotide supermatrix consisting of the concatenation of the 13 protein-coding genes and the two rRNA genes, (iii) a mitochondrial amino-acid supermatrix of the 13 protein-coding genes, and (iv) a mixed supermatrix of UCE nucleotides and mitochondrial amino-acid protein-coding genes. For all supermatrices, phylogenetic inference was performed with Maximum Likelihood (ML) as implemented in IQ-TREE v1.6.8 (Nguyen, Schmidt, von Haeseler, & Minh 2015) using a GTR+Γ_4_+I model for UCE and mitochondrial nucleotide supermatrices, a mtART+Γ_4_+I model partitioned by gene for mitochondrial amino acids matrices, and a partitioned model mixing a GTR+Γ_4_+I model for UCE nucleotides and a mtART+Γ_4_+I model for mitochondrial amino acids for the mixed supermatrices. Statistical node support was estimated using ultrafast bootstrap (UFBS) with 1000 replicates (Hoang, Chernomor, von Haeseler, Minh, & Vinh 2018). Nodes with UFBS values higher than 95% were considered as strongly supported. For all supermatrices, the congruence between the different topologies obtained with the four assemblers was evaluated by calculating quartet distances with Dquad (Ranwez, Criscuolo, & Douzery 2010).

## Results

### Assembly of UCE and mitochondrial datasets

*De novo* assembly of 501 UCE capture sequencing libraries was performed with four different assemblers: IDBA-UD, MEGAHIT, and MetaSPAdes via MitoFinder and Trinity via PHYLUCE. All assemblers provided different numbers of contigs (**Table 1**) ranging from 30,544 (IDBA-UD) to 114,392 (MEGAHIT) on average. The average computational time per assembly was highly variable among assemblers with Trinity being by far the slowest (35 CPUs, median total-time: 1h:06m:22s) and IDBA-UD the fastest (5 CPUs, median total-time: 0h:11m:01s), MEGAHIT (5 CPUs, median total-time: 0h:12m35s) being slightly slower, and MetaSPAdes (5 CPUs, median total-time: 0h:25m:44s) being about twice slower than the other two metagenomic assemblers (**Table 1 & Fig. 2A**).

**Table 1.**
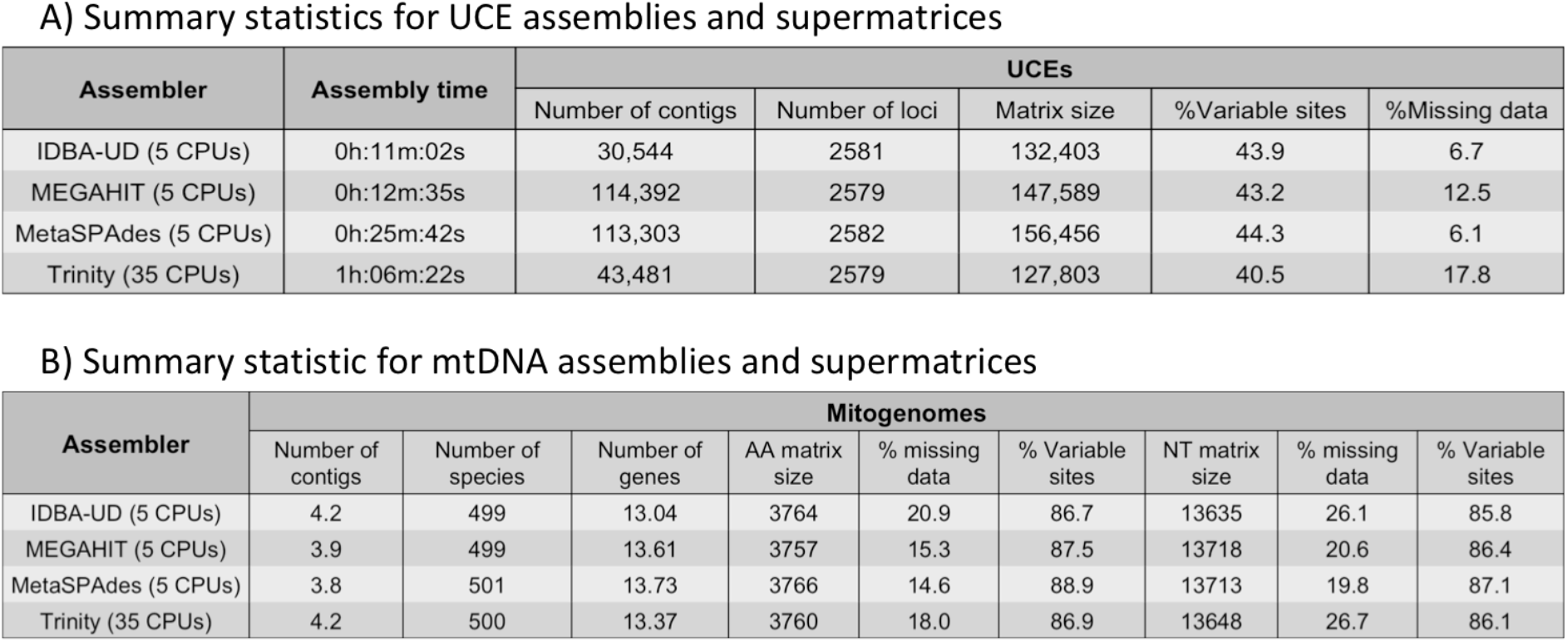
Summary statistics on assembly results according to the assembler used. The values are averages over the 501 assemblies, except for the assembly time, which is a median value. The two tables report specific statistics for A) ultraconserved elements data, and B) mitochondrial data. Note that 35 CPUs were used for Trinity whereas 5 CPUs were used for others assemblers.

A) Summary statistics for UCE assemblies and supermatrices
| Assembler | Assembly time | UCEs |  |  |  |  |
| --- | --- | --- | --- | --- | --- | --- |
|  |  | Number of contigs | Number of loci | Matrix size | %Variable sites | %Missing data |
| IDBA-UD (5 CPUs) | 0h:11m:02s | 30,544 | 2581 | 132,403 | 43.9 | 6.7 |
| MEGAHIT (5 CPUs) | 0h:12m:35s | 114,392 | 2579 | 147,589 | 43.2 | 12.5 |
| MetaSPAdes (5 CPUs) | 0h:25m:42s | 113,303 | 2582 | 156,456 | 44.3 | 6.1 |
| Trinity (35 CPUs) | 1h:06m:22s | 43,481 | 2579 | 127,803 | 40.5 | 17.8 |

**Table 1.** Summary statistics on assembly results according to the assembler used.
| Assembler | Mitogenomes |  |  |  |  |  |  |  |  |
| --- | --- | --- | --- | --- | --- | --- | --- | --- | --- |
|  | Number of contigs | Number of species | Number of genes | AA matrix size | % missing data | % Variable sites | NT matrix size | % missing data | % Variable sites |
| IDBA-UD (5 CPUs) | 4.2 | 499 | 13.04 | 3764 | 20.9 | 86.7 | 13635 | 26.1 | 85.8 |
| MEGAHIT (5 CPUs) | 3.9 | 499 | 13.61 | 3757 | 15.3 | 87.5 | 13718 | 20.6 | 86.4 |
| MetaSPAdes (5 CPUs) | 3.8 | 501 | 13.73 | 3766 | 14.6 | 88.9 | 13713 | 19.8 | 87.1 |
| Trinity (35 CPUs) | 4.2 | 500 | 13.37 | 3760 | 18.0 | 86.9 | 13648 | 26.7 | 86.1 |

**Figure 2.**
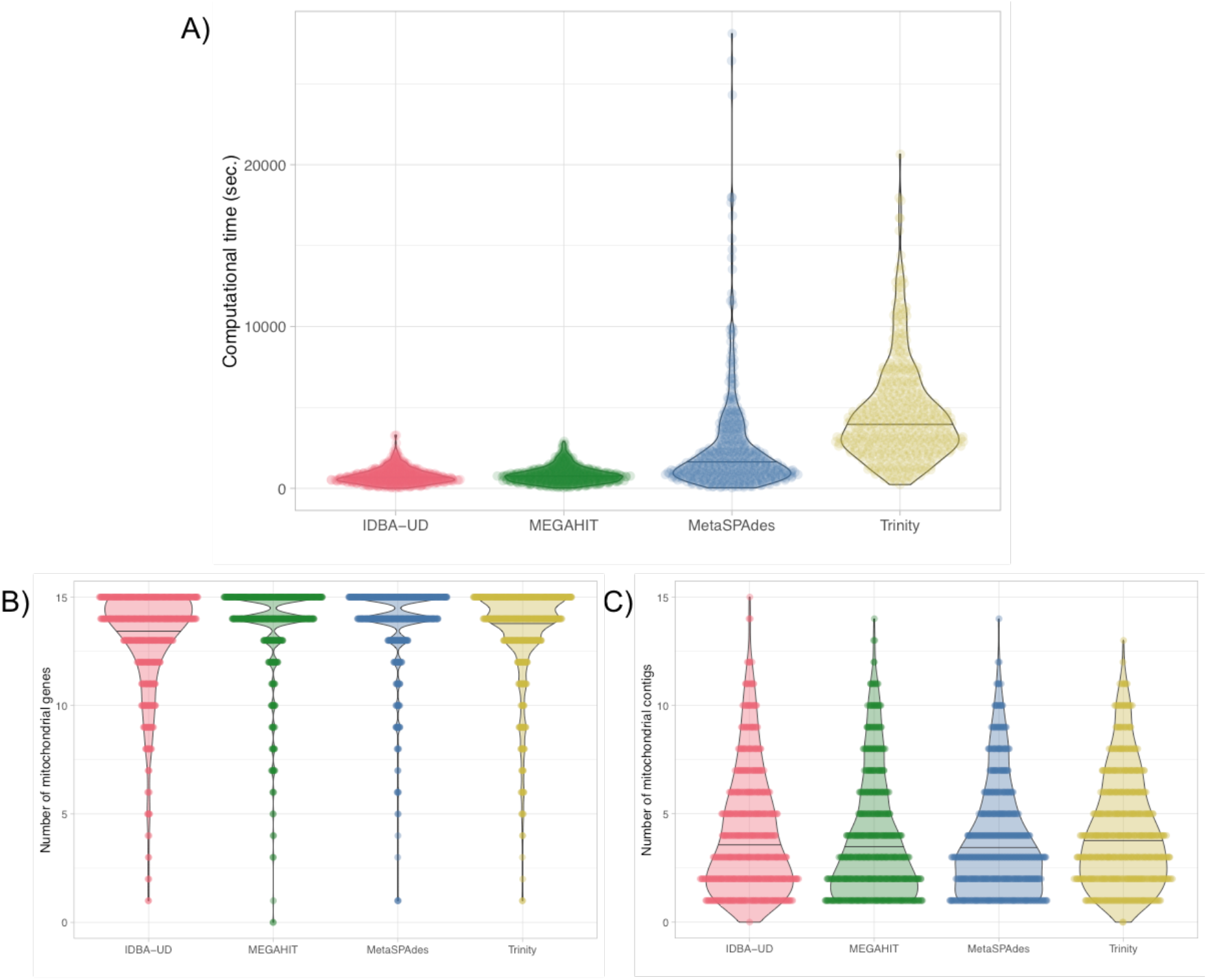
Comparison of the efficiency of the assemblers in terms of A) computational time, B) number of potentially mitochondrial contigs identified, and C) number of mitochondrial genes annotated. Violin plots reflect the data distribution with a horizontal line indicating the median. Note that for the three metagenomic assemblers, 5 CPUs were used compared to 35 CPUs for Trinity. Plots were obtained using PlotsOfData (Postma & Goedhart 2019).

The UCE supermatrices created by PHYLUCE for each of the four assemblers contained on average 2580/2590 UCE loci for Hymenoptera (**Table 1**). All matrices contained 501 species, but the size of the supermatrix and the percentage of missing data varied depending on the assembler (**Table 1**). Trinity, which is generally used as the default assembler in PHYLUCE, resulted in the shortest and most incomplete supermatrix with 2579 loci representing 127,803 sites (40.5% variable) and 17.8% missing data. Among metagenomic assemblers, MetaSPAdes provided the largest and most complete supermatrix with 2582 loci representing 156,456 sites (44.5% variable) and only 6.0% missing data. IDBA-UD retrieved 2581 loci representing 132,403 sites (43.9% variable) with only 6.7% missing data, and MEGAHIT resulted in a supermatrix with 2579 loci representing 147,589 sites (43.2% variable) but with 12.4% missing data. Note that less than 30 loci were retrieved for *Phalacromyrmex fugax* (between 4 and 27 loci depending on the assembler). This is congruent with the original publication in which this library was not included in phylogenetic analyses (Branstetter *et al.* 2017a). Accordingly, we removed the *Phalacromyrmex fugax* library (SRR5437956) from the dataset.

Depending on the assembler, mitochondrial signal was recovered in 499, 500, and 501 libraries out of a total of 501 (**Table 1**, **Fig. 2B)**. Overall, mitochondrial signal thus was detected in all libraries but only MetaSPAdes retrieved it in all species (**Appendix S2**). On average, 3.8 contigs per species was identified (**Table 1**, **Fig. 2B**) and 13.7 genes were annotated with MitoFinder (**Fig. 2C**). In 296/501 cases, MitoFinder was able to assemble a contig of more than 15,000 bp containing at least 13 annotated genes that likely represents the complete mitochondrial genome. In 52 of these cases, all 15 genes were annotated. In the remaining, the putative mitogenome contigs were missing one or two genes, mostly the short and divergent ATP8 (131/296), the 12S rRNA (29/296) and the 16S rRNA (10/296), which were present but not directly annotated by our blast-based procedure.

After alignment and cleaning, mitochondrial genes were used to create nucleotide and amino acid supermatrices. To be consistent with UCE analyses, and despite the recovery of some mitochondrial signal, we ignored *Phalacromyrmex fugax* in further analyses. In the nucleotide supermatrices (13 protein-coding + 12S and 16S rRNAs), we obtained 13 genes on average per species, which resulted in supermatrices with 13,679 nucleotide sites (86.4% variable) and 23.3% missing data on average (**Table 1**). In the amino acid matrices (13 protein-coding genes), we obtained supermatrices with 3762 amino acid sites (87.4% variable) and 17.2% missing data on average (**Table 1**).

### Barcoding analyses

A total of 534 COI sequences retrieved from the 501 MetaSPAdes assemblies were used to verify species identification of the UCE libraries (**Appendix S3**). In 42 cases, two or three COI sequence fragments were retrieved from the same UCE library. In seven of these cases, the slightly-overlapping COI fragments most likely resulted from bad assembly or erroneous annotation. However, in the 35 remaining cases, a genuine complete COI sequence overlapped with shorter fragments suggesting either cross-contaminations, nuclear mitochondrial DNA segments (NUMTs), endoparasites, or bacterial symbionts. For instance, in *Temnothorax* sp. mmp11 (SRR5809551), a 391 bp fragment annotated as COI by MitoFinder was found to be 98.2% identical to both the *Wolbachia pipientis* wAlbB and *Wolbachia* Pel strain wPip genomes, which are bacterial endosymbionts of the mosquitoes *Aedes albopictus* and *Culex quinquefasciatus*, respectively. Also, in *Sericomyrmex bondari* (SRR5044901) and *Sericomyrmex mayri* (SRR5044856) short COI fragments best matched with nematodes. However, in the 312 cases for which COI barcoding allowed to confirm the species identity of the UCE libraries, we did not detect any obvious cases of cross-contaminations where the COI extracted from a given library would have been identical to the one of another library (**Appendix S3**).

### Phylogenetic results

The ML topologies inferred from the different UCE supermatrices were very similar with an average quartet distance of 0.005 among assemblers (**Appendix S4**). However, the percentage of supported nodes (UFBS > 95) differed depending on the assembler: IDBA-UD (91.37%), MetaSPAdes (89.96%), MEGAHIT (89.56%), and Trinity (85.85%). In the following, we only discuss the phylogenetic results obtained with MetaSPAdes that provides the most comprehensive assemblies for both UCE and mitochondrial data (**Table 1**). The following 12 well-established subfamilies were retrieved with maximal UFBS support (100%): Aneuretinae, Amblyoponinae, Dolichoderinae, Dorylinae, Ectatomminae, Formicinae, Heteroponerinae, Myrmeciinae, Myrmicinae, Pseudomyrmecinae, and Ponerinae (**Fig. 3A**). The two supergroups Formicoid and Poneroid were also retrieved with maximal UFBS support, as well as consensual phylogenetic relationships among Formicoid subfamilies (Ward 2014).

**Figure 3.**
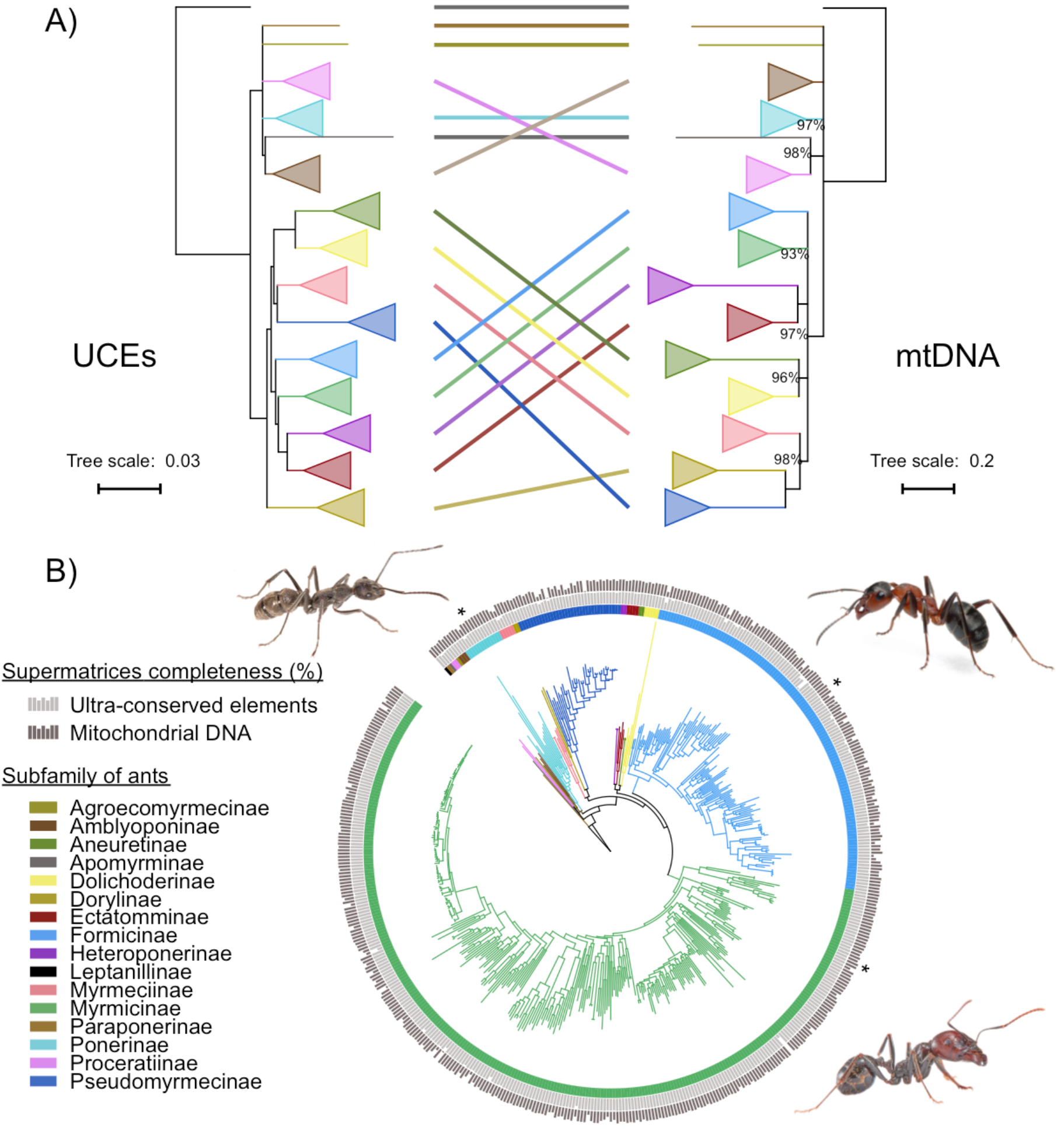
Phylogenomic relationships of Formicidae. A) The topology reflects the results of phylogenetic analyses based on UCEs amino acid mitochondrial supermatrix. Histograms reflect the percent of UCE (dark grey) and mitochondrial genes (light grey) recovered for each species. Illustrative pictures (*): *Diacamma* sp. (Ponerinae; top left), *Formica* sp. (Formicinae; top right), and *Messor barbarus* (Myrmicinae; bottom right). B) Mito-nuclear phylogenetic differences for ancient relationships. Clade corresponding to subfamilies were collapsed. Inter-subfamily relationships with UFBS < 95 were collapsed. Non-maximal node support values are reported.

For mitochondrial matrices, the percentage of supported nodes (UFBS> 95)with nucleotides also differed depending on the assembler and was higher than with the amino acids: MetaSPAdes (84.5% *vs.* 80.1%), MEGAHIT (84.0% *vs.* 79.4%), Trinity (83.3% *vs.* 80.4%), and IDBA-UD (80.2% *vs.* 78.0%). However, ML mitogenomic trees inferred from amino acids were more congruent with UCE topologies than the ones inferred from the mitochondrial nucleotides (average quartet distance = 0.035 *v.s.* 0.063; **Appendix S4**). Among assemblers, the ML topologies inferred with amino acid matrices were highly congruent with an average quartet distance of 0.007 (**Appendix S4**). In the ML tree obtained with the MetaSPAdes supermatrix (**Fig. 3B**), all ant subfamilies were retrieved with maximal UFBS support values except for Myrmicinae (93%), Ponerinae (97%), and Proceratinae (99%) (**Fig. 3A**). However, relationships among subfamilies were not congruent with UCE phylogenomic inferences except for Heteroponerinae + Ectatomminae (UFBS = 100) and Dolichoderinae + Aneuretinae (UFBS = 96) (**Fig. 3A**).

Finally, phylogenetic inference carried on mixed supermatrices composed of UCEs and mitochondrial amino acids resulted in ML topologies that were also highly similar among assemblers with an average quartet distance of 0.006 (**Appendix S4**). The percentage of supported nodes (UFBS > 95) were: IDBA-UD (91.2%), MEGAHIT (92.8%), MetaSPAdes (92.2%), and Trinity (90.4%). As with UCE matrices, the 12 well-established subfamilies, the two supergroups Formicoid and Poneroid and consensual Formicoid inter-subfamilies relationships (Ward 2014) were all retrieved with maximal UFBS support.

## Discussion

### Metagenomic assemblers are efficient tools for assembling UCEs

Currently, genomic and transcriptomic *de novo* assemblers are commonly used to assemble UCE loci from DNA capture sequencing data (Faircloth 2016). Since metagenomic assemblers such as IDBA-UD, MEGAHIT, and MetaSPAdes have been designed to account for variance in sequencing coverage, they seem to be well adapted for targeted enrichment or DNA sequence capture data. Our results show that metagenomic assemblers are indeed more efficient at assembling UCE loci than the classically used, but computationally intensive, Trinity transcriptomic assembler. As a consequence, they could lead to datasets containing more variable sites, less missing data, and increased phylogenetic signal (**Table1**). Indeed, the topologies obtained with the metagenomic assemblers are very similar to the topology obtained with the Trinity-based supermatrix, contain a higher number of supported nodes (UFBS ≥ 95%), and are consistent to previous studies (Ward 2014). Furthermore, assemblies obtained with the three metagenomic assemblers provide variable numbers of contigs (ranging from 30,544 to 114,392) resulting in differences in the completeness of the matrices (6.0% to 17.8% of missing data for UCE matrices and 29.9% to 41.3% for mitochondrial matrices) and in numbers of variable sites (for UCE, 40.5% to 44.3%; for mtDNA, 77.2% to 79.0%). Interestingly, for both UCE matrices and mtDNA matrices, MetaSPAdes consistently provides more loci, more variable sites, and less missing data. In addition, mitochondrial signal was extracted from all libraries only using MetaSPAdes within Mitofinder. Despite a computation time on average twice that of the other two metagenomic assemblers, MetaSPAdes is the more efficient assembler for ant UCEs. This software therefore provides a much needed alternative to Trinity for efficiently assembling hundreds of UCE libraries.

### Mitochondrial signal can systematically be extracted from UCE capture data

Ultraconserved elements are key loci exploited as target capture sequences in an increasing number of phylogenomic studies. DNA sequence capture methods are used to efficiently enrich targeted DNA regions in library preparation prior to sequencing, but non-targeted regions are always sequenced in the process resulting in so called “off-target reads”. Interestingly, off-target reads could represent up to 40% of the sequenced reads in exome capture experiments (Chilamakuri *et al.* 2014) and many contigs not belonging to targeted UCE loci are typically assembled from UCE capture data (e.g. Smith, Harvey, Faircloth, Glenn, & Brumfield, 2014; Faircloth *et al.* 2015). Given this high proportion of off-target reads, we can expect that mitochondrial DNA could be found as off-target sequences in many target enrichment data. Accordingly, several studies have succeeded in extracting mtDNA from UCE libraries (e.g. Smith *et al.* 2014; do Amaro *et al.* 2015). The development of MitoFinder allowed the automatic extraction of mitochondrial signal from all 501 ant UCE libraries. This maximum success rate indicates that this approach is highly efficient at least in Formicidae. However, the success in retrieving mitochondrial sequences, ultimately depends on the number of mitochondria contained in the tissue used for DNA extraction and library preparation. As expected, mitochondrial off-target reads are much more common in muscle and heart than in lung tissues in human (D’Erchia *et al.* 2015). Similarly, mitochondrial sequences are probably rare or absent in library constructed from vertebrate blood, even in birds in which nucleated red blood cells contain mitochondria, but in very low numbers (Reverter *et al.* 2016). In invertebrates, our case study with 100% success rate in ant UCEs demonstrates that mitochondrial sequences could probably be easily retrieved for many arthropod taxa as a by product of target enrichment sequencing experiments.

### The value of complementary mitochondrial signal

Mitochondrial sequences could provide interesting and important complementary information compared to nuclear sequences. First, mtDNA can be used to confirm the identity of the species sequenced for conserved UCE loci. Here, we were able to confirm the identification of 312 ant species out of the 501 UCE libraries using COI barcoding without revealing a single case of obvious species misidentification. Given that ant UCE libraries have been constructed from museum specimens, the 501 COI sequences we annotated could be used as reference barcoding sequences in future studies. Then, even though we did not detect such cases, the high mutation rate and the absence of heterozygous sites in mtDNA also make it well adapted for cross-contamination detection analyses (Ballenghien *et al.* 2017).

Nevertheless, mitochondrial markers also have some well identified limitations (Galtier *et al.* 2009). First, mtDNA could be inserted in the nuclear genome in the form of NUMTs (Bensasson, Zhang, Hartl, & Hewitt 2001). NUMTs could potentially be assembled as off-target contigs in DNA capture libraries and we might have indeed extracted some fragments corresponding to NUMTs for the COI gene using MitoFinder (**Appendix S2**). Theoretically, NUMTs could be picked up by analysing the coverage of putative mitochondrial contigs as they are expected to have a coverage comparable to other off-targets nuclear contigs, whereas genuine mitochondrial contigs should have a higher coverage. A second limitation of mtDNA exists in arthropods where maternally inherited intra-cellular bacteria are frequent. Among those bacteria, *Wolbachia* is particularly widespread and could distort the mitochondrial genealogy when a particular strain spreads within the host species hitchhiking its linked mitochondrial haplotype (Cariou, Duret, & Charlat 2017). *Wolbachia* is frequent among ants and could therefore be responsible of some mito-nuclear discordance (Wenseleers *et al.* 1998). We indeed discovered such an instance with a *Wolbachia* COI sequence identified in *Temnothorax* sp. mmp11 (SRR5809551), which was confirmed by several assembled contigs matching to *Wolbachia* strain genomes in this sample.

Beyond the methodological aspects of species identification and potential cross-contamination detection, mitochondrial sequences could also be useful to tackle fundamental evolutionary questions. UCEs have also proved to be useful genetic markers for phylogeography and for resolving shallow phylogenetic relationships (Musher & Cracraft 2018; Smith *et al.* 2014). In this context, mtDNA could also bring complementary information. In most animals, mtDNA has a maternal inheritance without recombination, which means that all mitochondrial genes behave as a single locus. This simplifies the interpretation of the phylogenetic pattern between closely related species or within subdivided populations of a species. Mito-nuclear phylogenetic discordance could also reveal interesting phenomena involving hybridization, sex-biased dispersal, and introgression (Toews & Brelsford, 2012). In practise, hybridization events are often identified using mito-nuclear discordance (Li *et al* 2016) and in some cases, the mitochondrial introgression events have proven to be adaptive (Seixas, Boursot & Melo-Ferreira 2018). Nevertheless, in our ant case study, a detailed comparison of mitochondrial and UCE phylogenies did not allow revealing convincing occurrences of such discordances.

### Ant phylogenetic relationships from 500 UCE and mitochondrial data

Both nuclear and mitochondrial data retrieved the most consensual phylogenetic relationships in the ant phylogeny (Ward 2014; Branstetter *et al.* 2017b; Borowiec *et al.* 2019). Twelve Formicidae subfamilies were recovered as monophyletic in all analyses, both with the nuclear and mitochondrial datasets, confirming their robustness. However, the well-defined inter-subfamily relationships within Formicoids (Ward 2014; Branstetter *et al.* 2017; Borrowiec *et al.* 2019) were only supported by the UCE dataset, but not by the mitochondrial amino acid dataset. For example, the army ant subfamily (Dorylinae) was not retrieved as the sister-group of all other Formicoids, but was the closest relative of Pseudomyrmicinae (UFBS = 100). Similarly, contradicting the classical and well-defined relationship of Heteroponerinae+Ectatomminae as the sister-group of Myrmicinae (Ward 2014; Branstetter *et al.* 2017; Borrowiec *et al.* 2019), the mitochondrial dataset supported an alternative relationship with Dolichoderinae+Aneuretinae (UFBS = 96). These differences suggest that mitochondrial data might be not well-suited to resolve ancient phylogenetic relationships at the ant inter-subfamily level, even if they look suitable for more recent nodes such as intra-subfamily relationships.

Interestingly, these topological incongruences between UCEs and mitochondrial genes also featured different topologies regarding the existence of the Poneroid taxa, a controversial clade not always retrieved depending on the studies (Ward 2014), but that tends to be retrieved in the most recent studies (Branstetter *et al.* 2017; Borrowiec *et al.* 2019; UCE dataset in this study) and is not recovered by our mitochondrial amino acid dataset (**Fig. 3B**). The same applies to the phylogenetic placement of Apomyrminae, a subfamily either grouped with Leptanillinae or Amblyoponinae in past studies (Ward 2014), but that was grouped with Proceratinae in our mitochondrial dataset (UFBS = 98; **Fig. 3B**). For such controversial nodes, our study demonstrates that the nature of the phylogenetic markers can provide different results. Such differences between nuclear and mitochondrial data might be due to the substitutional saturation of mitochondrial data even at the amino acid level. This problem may actually be exacerbated in hymenopteran mitochondria that possess high AT content translating into strongly biased codon usage potentially leading to phylogenetic reconstruction artefacts (Foster, Jermiin & Hickey 1997; Foster & Hickey 1999). Interestingly, such differences between mitochondrial and nuclear inference for ancient phylogenetic relationships, is not observed with insects with less AT-rich mitochondrial genomes such as, for instance, swallowtail butterflies (Condamine, Nabholz, Clamens, Dupuis & Sperling, 2018; Allio *et al.* 2019) or tiger beetles (Vogler & Pearson 1996). This calls for additional studies on both controversial and consensual ant inter-subfamily relationships with more comprehensive genome-wide datasets.

## Conclusions

In this study, we developed the MitoFinder tool to automatically extract and annotate mitogenomic data from raw sequencing data in an efficient way. For the assembly step of our pipeline, we tested four different assemblers and showed that MetaSPAdes is the most efficient and accurate assembler for both UCE and mitochondrial data. Applying MitoFinder to ants, we were able to extract mitochondrial signal from 501 UCE libraries. This demonstrates that mitochondrial DNA can be found as off-target sequences in UCEs sequencing data. Interestingly, mitochondrial DNA extracted from UCE libraries can also be used to: (i) confirm species identification with barcoding methods, (ii) highlight potential sample cross-contamination, and (iii) reveal potential cases of mito-nuclear discordance caused by hybridization events leading to mitochondrial introgression. Finally, MitoFinder was developed with UCE libraries but our approach should also work with data obtained from other capture methods in which numerous off-targets reads are sequenced, as well as with transcriptomic and whole genome sequencing data, in which mitochondrial reads are overrepresented.

## Acknowledgements

This paper is dedicated to the memory of graduate student Alex Schomaker-Bastos (1992-2015) who was assassinated by the time he was writing the mitoMaker program on which we built upon for the annotation module of MitoFinder. We also thank Fabien Condamine for providing helpful comments on a previous version of the manuscript. This work has been supported by grants from Investissements d’Avenir of the Agence Nationale de la Recherche (CEBA: ANR-10-LABX-25-01; CEMEB: ANR-10-LABX-0004), and the European Research Council (ERC-2015-CoG-683257 project ConvergeAnt). This is contribution ISEM 2019-XXX of the Institut des Sciences de l’Evolution de Montpellier.

## Data accessibility

The MitoFinder software is available from XXXXXX. Annotated mitogenomes and partial mitogenomic contigs have been deposited in GenBank (Accession Numbers XXXXX-XXXXX). The full analytical pipeline, phylogenetic datasets and corresponding trees can be retrieved from zenodo.org (DOI:10.5281/zenodo.3231390).

## Authors’ contributions

RA and FD conceived the ideas and designed methodology, analysed the data, and led the writing of the manuscript; RA implemented the MitoFinder software in part using code previously written by AS-B; JR, FP, and BN contributed to the writing of the manuscript. All authors contributed critically to the drafts and gave final approval for publication.

## Supporting information

**Appendix S1.** List of the 501 UCE libraries (SRA accessions) and associated metadata.

**Appendix S2.** Summary statistics on mitochondrial signal recovered per species and depending on the assembler used. The table provides the number of contigs and genes recovered with MitoFinder and the size of each annotated gene.

**Appendix S3.** Summary statistics of barcoding analyses. Detailed results for both BOLDsystem and Megablast analyses are provided for each COX1 recovered with MitoFinder using MetaSPAdes.

**Appendix S4.** Detailed results of tree distance analyses realized with Dquad (Ranwez, Criscuolo, & Douzery 2010). Trees obtained with each assembler with mitochondrial amino acid supermatrix, mitochondrial nucleotide supermatrix, and UCE nucleotide supermatrix were compared with each others.

